# Early-stage idiopathic pulmonary fibrosis is characterized by bronchoalveolar accumulation of SPP1^+^ macrophages

**DOI:** 10.1101/2023.12.06.569201

**Authors:** Jiangyan Yu, Jake Thomas, Jessica Haub, Carmen Pizarro, Miaomiao Zhang, Leonie Biener, Matthias Becker, Lili Zhang, Theodore S. Kapellos, Wolfgang Schulte, Joachim L. Schultze, Jan Hasenauer, Dirk Skowasch, Andreas Schlitzer

## Abstract

Patients affected by idiopathic pulmonary fibrosis (IPF), a progressive chronic and eventually fatal lung disease with unknown cause, suffer from delayed diagnosis and limited personalized treatment options due to the lack of predictive and staging relevant disease markers. Prior studies focused on the cellular and transcriptomic changes found in lung tissue during terminal IPF-associated lung fibrosis. In clinical routine, bronchoscopy is applied for diagnosis and staging of suspected IPF affected individuals, thus providing us with a clinically applicable window, to study the cellular and transcriptional changes within affected individuals at the time of diagnosis. Here we study a cohort consisting of 11 IPF affected individuals and 11 healthy controls. We investigate their single cell transcriptomic profile alongside the surface phenotype of alveolar-resident immune cells. Single cell transcriptional analysis reveals accumulation of SPP1^+^ macrophages (SPP1^+^Mφ) within the alveolar space during early stage diagnosed IPF, in the absence of a decline in lung function. SPP1^+^Mφ were characterized by high expression of lipid storage and handling genes, accumulation of intracellular cholesterol and expression of proinflammatory cytokine expression. *In silico* developmental trajectory analysis revealed that SPP1^+^Mφ are likely derived from classical monocytes. Finally, to confirm the clinical diagnostic value of SPP1^+^ Mφ, we developed a flow cytometry panel to rapidly identify and test for the presence of SPP1^+^ Mφ in bronchoalveolar lavage samples. The new panel has confirmed the increased frequency of SPP1^+^ Mφ in early-IPF in an independent validation cohort. Taken together, we show that SPP1^+^ Mφ are associated with early IPF pathogenesis in the absence of a decline in lung function, thus providing a clinically valuable markers for diagnosis of early IPF in clinical practice.

**Highlights:**

1. IPF patients can be stratified into early and advanced disease groups based on clinical features.
2. Single cell transcriptomic and high dimensional flow cytometry enabled mapping of the myeloid compartment across the continuum of idiopathic pulmonary fibrosis associated disease states.
3. SPP1^+^Mφ accumulated during early clinical stage of IPF.
4. Development of a clinically applicable SPP1⁺Mφ identification panel.

## Introduction

Idiopathic pulmonary fibrosis (IPF) is a rare progressive fibrotic lung disease with limited treatment options and a median survival of 2-5 years after definite diagnosis (Kaunisto et al., 2019; Lederer and Martinez, 2018). Due to the poor understanding of disease pathogenesis and lack of diagnostic molecular markers, diagnosis of IPF often follows an exclusion process of other known causes of lung fibrosis (Martinez et al., 2017; Raghu et al., 2018). Despite recent advances in clinical imaging and multidisciplinary evaluation of cases, median time to diagnosis is 2 years, leading to a considerable delay in the start of treatment significantly lowering patient survival (Hoyer et al., 2019; Martinez et al., 2017). Thus, a high clinical need exists for the identification of cellular and molecular markers useful for early IPF diagnosis.

Recent single-cell transcriptional analysis of end-stage IPF-associated lung transplant tissue has revolutionized our understanding of the molecular pathogenesis of IPF (Adams et al., 2020; Carraro et al., 2020; Habermann et al., 2020; Xu et al., 2017). These studies detailed both immune and non-immune compartments and revealed an unappreciated macrophage heterogeneity within fibrotic foci of affected lungs. Further single-cell transcriptomic studies could shed further light on the interplay of non-immune cells, such as KRT17^+^ basal cells as drivers of myofibroblast differentiation and proliferation (Jaeger et al., 2022). Additionally mouse models have indicated macrophages as major regulators of fibrotic disease (Ucero et al., 2019; Zhang et al., 2020). Similarly in terminal lung tissue samples of IPF affected individuals profibrotic macrophages, marked by the expression of *MERTK* and *SPP1* were found indicating a possible disease driving or modifying role in human disease (Adams et al., 2020; Morse et al., 2019; Reyfman et al., 2019). However, diagnostic value and the immediate benefit to patients of these findings remains low as lung tissue biopsy material is not obtained in routine clinical practice anymore (Hutchinson et al., 2015).

Bronchoscopy is a minimally invasive procedure to flush the lung with saline solution via a bronchoscope, and is used to diagnose IPF in clinical routine (Pesci et al., 2010). Bronchoscopy obtained patient material has recently been used to identify CCR5, CCR6, CXCR3 and CXCR5 as biomarkers for transplant-stage IPF (Sivakumar et al., 2019). Additionally, broncho-alveolar resident CD71^+^ macrophages were identified to be enriched in IPF-affected individuals but not in control subjects (Allden et al., 2019). Thus, analysis of bronchoalveolar lavage samples yields tangible and clinically applicable results to further understand IPF pathogenesis.

Here, using cellular specimens obtained from IPF or control patients we characterize the transcriptomic and phenotypic landscape of broncho-alveolar resident mononuclear phagocytes and link the abundance of SPP1^+^ macrophages (SPP1^+^Mφ) to an early IPF-disease stage which presents clinically without any obvious lung function decline. Further analyses of SPP1^+^Mφ revealed upregulated lipid handling and storage pathways alongside the accumulation of intracellular cholesterol. Transcriptional developmental trajectory analysis suggests a monocyte-to-macrophage differentiation trajectory within the bronchoalveolar space. Lastly, we develop and test a clinically applicable flow cytometry panel confirming the association of SPP1^+^Mφ with early IPF pathogenesis in an independent cohort. Collectively we show that accumulation of lipid-laden bronchoalveolar-resident monocyte-derived SPP1^+^Mφ is a clinically tangible feature of early stage IPF, enabling a faster diagnosis and treatment onset in at-risk patients.

## Results

### Single-cell transcriptomic analysis determined 8 distinct types of macrophages in the human alveolar space

Clinically newly diagnosed IPF patients are not staged as per their disease progression due to the lack stratification criteria informing disease management. However, lack of knowledge related to disease progression leading to terminal fibrotic outcomes hamper our understanding of available treatment windows in early patients. To determine IPF stage specific cellular states and according molecular markers, we stratified IPF patients into subgroups using available clinical parameters. To achieve this, we classified 12 subjects with chronic cough but without any sign of fibrosis in the lung (hereafter referred to as “control”) and 50 IPF patients, using unsupervised hierarchical clustering of all available lung function parameters (**Figure 1A, Table S1**). After exclusion of highly correlated parameters (**Figure S1A**), six lung function parameters were subjected for cluster analysis, yielding two major clusters of patients: cluster-I contained mostly IPF patients while cluster-II consisted of IPF patients and controls (**Figure 1B**). In order to understand why cluster-I showed a mixed phenotype between control and IPF-affected patients, we realized a strong correlation between cluster identity and FEV1 and DLco predicted values (**Figure 1C-D**), suggesting that IPF patients within cluster-I displayed a similar degree of lung function as unaffected control individuals. Taken together these findings prompted us to define IPF affected individuals within cluster-I as early-IPF, whereas patients in cluster-II, showing a strong decline in FEV1 and DLco, as advanced-IPF group. This lung-function based stratification was further consistent with the level of lung fibrosis as measured by high-resolution computed tomography (HRCT) imaging data, which indicated that patients in the advanced-IPF group had significantly higher proportions of reticular and areas of ground glass opacity within their lungs as compared to early-IPF group individuals, indicating more advanced fibrosis (**Figure S1B**). Collectively using these stratification criteria we reveal early and advanced IPF phenotypes within our cohort.

**Figure 1.**
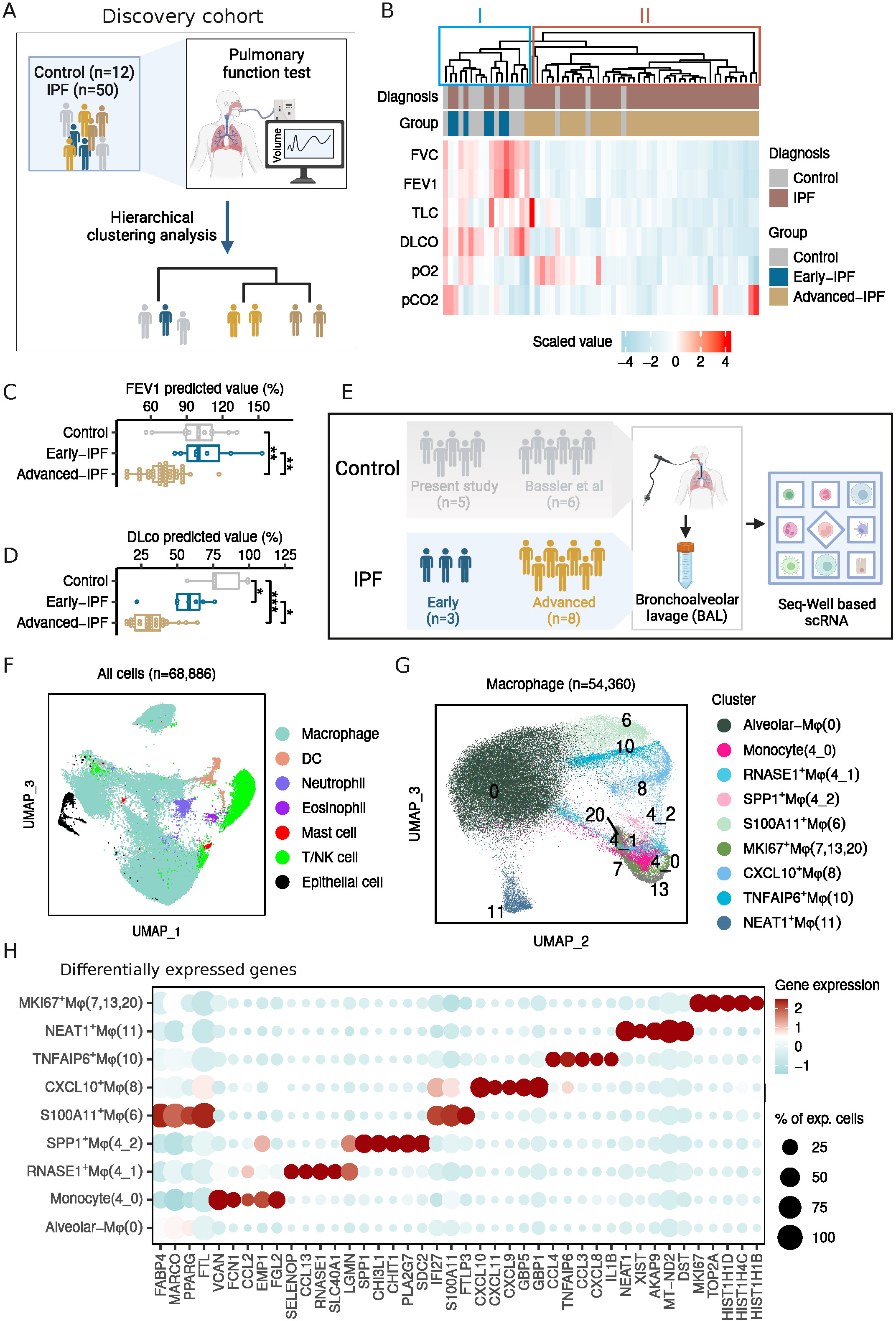
Single cell transcriptomic analysis revealed macrophage heterogeneity in the bronchoalveolar space. (A) Schematic workflow of the cluster analysis to stratify IPF patients. (B) Heatmap of lung function parameters of patients. FVC: forced vital capacity; FEV_1_: forced expiratory volume in 1 second; TLC: total lung capacity; D_LCO_: carbon monoxide diffusion capacity; pO2: partial pressure of oxygen; pCO2: partial pressure of carbon dioxide. (C) Percentage of predicted FEV_1_ value in controls and two IPF groups. (D) Percentage of predicted D_LCO_ value in controls and two IPF groups. (E) Schematic workflow of Seq-Well based single-cell sequencing analysis. (F) Major cell populations determined in BALF samples. (G) UMAP embedding of macrophage subpopulations. (H) Top 5 feature genes for each macrophage subpopulation by DEG analysis.

To link clinical parameter enabled patient stratification to cellular and molecular phenotypes we probed the cellular composition of the according BALF samples of early and advanced IPF group patients (n=19) using high-dimensional flow cytometry (**Figure S1C**). We observed that the human bronchoalveolar space is mainly composed of myeloid cells (**Figure S1D**). Within myeloid cells CD169^+^CD206^+^ alveolar macrophages dominated the human bronchoalveolar space (64.3% - 96.5%, **Figure S1E)**, in line with previous reports (Baßler et al., 2022; Yu et al., 2016). Furthermore, we found that advanced-IPF group individuals has significantly increased abundance of eosinophils (*p* < 0.01) and DC1 (*p* < 0.05) as compared to patients in the early-IPF group or the control group (**Figure S1E**), suggesting these cells might play a central role during the later stages of IPF progression.

Next to further understand early and advance-IPF associated transcriptomic programs and to provide an unbiased characterisation of cell types in the bronchoalveolar space we performed single-cell RNA sequencing (scRNA-seq) analysis broncho-alveolar associated cells in 11 IPF patients and 5 controls. To increase the statistical power, we integrated our scRNA data with a recently published dataset that contains 6 BALF samples from controls (**Figure 1E**) (Baßler et al., 2022). Cluster analysis determined a total of 20 distinct cell populations including monocytes/macrophages (9 clusters), dendritic cells (2 clusters), T/NK cells (3 clusters), granulocytes (3 clusters) and epithelial cells (3 clusters, **Figure 1F, Figure S2A-B)**, annotations of which were consistent with previous report (**Figure S2C-D**) (Travaglini et al., 2020). In line with our flow cytometry data, scRNA analysis revealed that macrophages were the most abundant cells in the human alveolar space (**Figure 1F**, **Figure S2E**). Re-clustering of the monocytes/macrophage cluster at an increased resolution yielded 9 sub-clusters, including one cluster of monocytes and 8 transcriptionally distinct macrophage subpopulations in BALF samples (**Figure 1G**).

To assess the heterogeneity of macrophages we performed differentially expressed gene (DEG) analysis (**Figure 1H, Table S2**). Cells in cluster 0 expressed classical alveolar macrophage genes including *FABP4* and *FTL* (**Figure S2D**). Three clusters (7, 13 and 20) were characterized by proliferating-associated genes including *TOP2A*, *MKI67* and *NUSAP1* as well as histone-related genes such as *HIST1H1D*, thus we grouped these three cluster into one population as MKI67^+^Mφ (**Figure S2D**). Cluster 4_1 (RNASE1^+^Mφ) was defined by *RNASE1* and *SELENOP* expression, resembling interstitial macrophages typically found within the lung parenchyma (Chakarov et al., 2019; Schyns et al., 2019), while cluster 4_2 (SPP1^+^Mφ) highly expressed *SPP1*, a gene that encodes for osteopontin, an integrin-binding glyco-phosphoprotein. *SPP1*-expressing macrophages have been previously described in a range of chronic fibrotic diseases and within the tumour microenvironment (Fabre et al., 2023; Ramachandran et al., 2019). Furthermore, DEG analysis revealed four types of activated macrophage subpopulations. Cluster 6 (S100A11^+^Mφ) was highly enriched for *S100A11* together with genes related to immune response activation signalling pathways (*C1QC*, *LGALS3*, *TYROBP* and *FCER1G*). Cluster 8 (CXCL10^+^Mφ) featured genes related to interferon γ response (*CXCL10*, *CXCL9*, *IFIT2* and *IFIT3*). Cluster 10 (TNFAIP6^+^Mφ) exhibited strong expression of genes related to inflammatory response including *TNFAIP6*, *CCL3* and *CCL4* (Mittal et al., 2016). Cluster 11 (NEAT1^+^Mφ) was characterized by expression of *NEAT1*, a lncRNA previously linked to regulation of the macrophage inflammatory response (Zhang et al., 2019).

Together these analyses revealed a highly diverse macrophage ecosystem within the human bronchoalveolar space from healthy and IPF patients.

### SPP1**^+^**M**φ** accumulate in the broncho-alveolar space of early stage IPF patients

With the cell populations determined from whole transcriptome analysis, we next sought to assess disease state related cell types. In line with the afore described flow cytometry results we observed that macrophages are the most redundant population in the bronchoalveolar space (**Figure S3A**, **Table S4**). Given the remarkable heterogeneity of macrophages within the human alveolar space, we investigated whether any macrophage subpopulations were related to IPF progression (**Figure 2A-B**). Three subpopulations (RNASE1^+^Mφ, TNFAIP^+^67Mφ and MKI^+^67Mφ) were found to be enriched in the advanced-IPF group, but this difference might be due to the low frequency in control samples from the published dataset (**Figure 2B**, **Figure S3B**). Nevertheless, quantification the frequency of macrophage subpopulations out of all macrophages revealed a predominant enrichment of SPP1^+^Mφ in the early-IPF group, and this difference is irrelevant to the source of control samples (**Figure 2C**, **Figure S3B**). Together this data has revealed that SPP1^+^Mφ were predominantly enriched in early-stage IPF patients.

**Figure 2.**
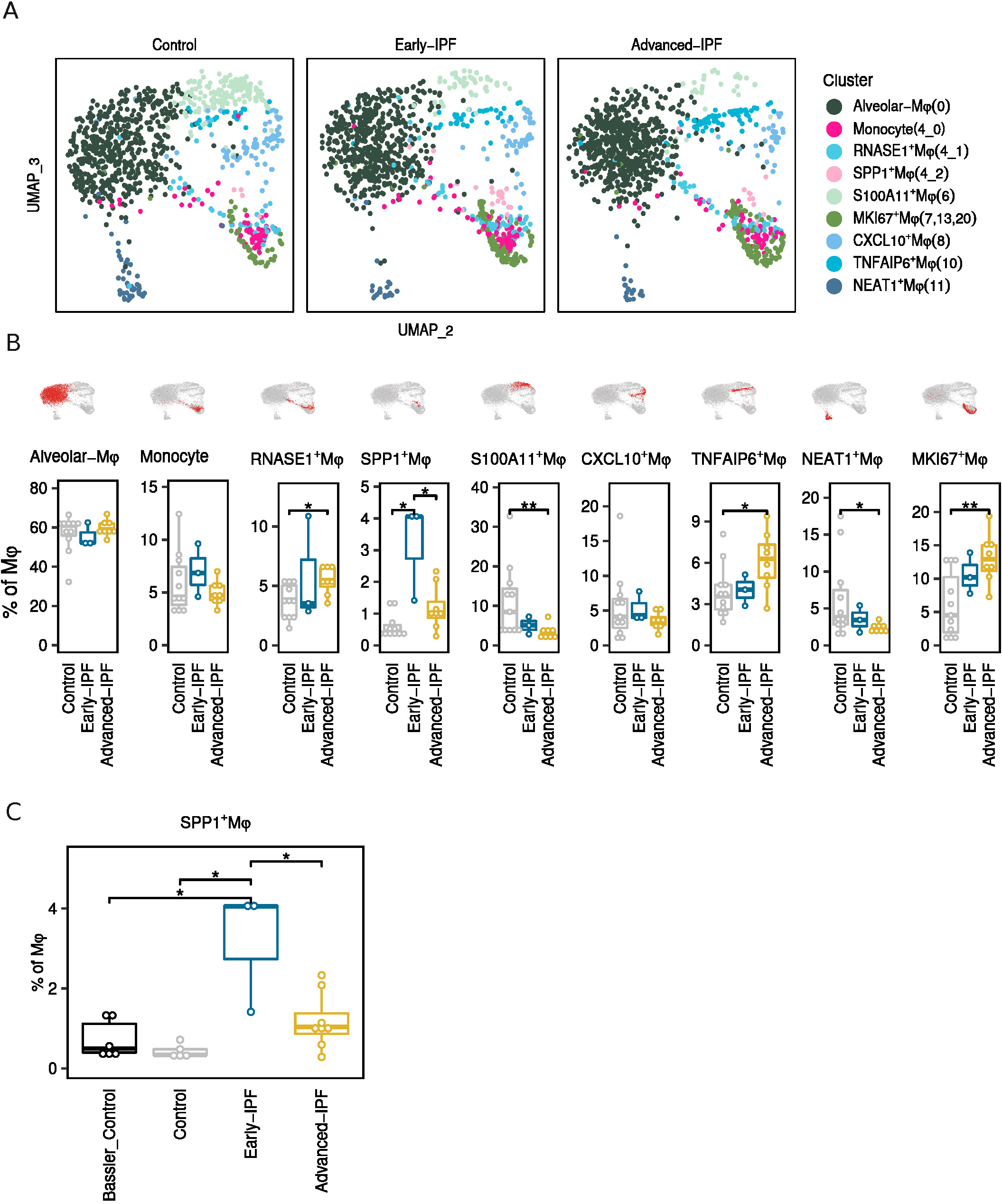
SPP1^+^Mφ were predominantly accumulated at the early stage of IPF development. (A) UMAP embedding of macrophage subpopulations separated by disease groups. To have an intuitive view of cell composition in each group, same number of cells (n=1000) were shown in each group. (B) Frequency of each subpopulation in the control and two disease groups. \**p* < 0.05, \*\**p* < 0.01, Wilcox-test. (C) Frequency of SPP1^+^Mφ in the control and disease groups. \**p* < 0.05, Wilcox-test.

### SPP1**^+^**M**φ** were characterized by genes related to lipid metabolism and are likely derived from monocytes upon fatty acid stimulation

Previous studies in mice have shown that macrophages in the lung are comprised of two ontogenically distinct populations: tissue-resident macrophages seeded during embryonic development and macrophages derived from adult blood monocytes (Guilliams et al., 2013; Mass et al., 2016; Schyns et al., 2019). Therefore, we sought to assess the origin of macrophage subpopulations identified in our study. To achieve this we made use of the MoMac-VERSE database that integrated mononuclear phagocytes from 13 human tissues and determined monocyte-derived macrophages and tissue-resident macrophages (**Figure 3A)** (Mulder et al., 2021). After transferring cluster labels from MoMac database to our macrophage subpopulations, we observed a vast majority of the macrophages were predicted to be tissue-resident macrophages, consistent with their identity as alveolar macrophages (**Figure 3B**). Meanwhile, we observed a strong transcriptional similarity between BALF SPP1^+^Mφ and monocyte-derived TREM2^+^Mφ from the MoMac database (**Figure 3B**, **Figure S4A**). To assess whether SPP1^+^Mφ are derived from monocytes, we performed trajectory analysis by embedding all BALF classical monocytes and monocyte-derived macrophages on the diffusion map also confirmed the differentiation process of monocytes to SPP1^+^Mφ (**Figure 3C**). Along the pseudotime developing from monocyte to SPP1^+^Mφ, genes related to monocyte identity including *VCAN*, *FCN1* and *S100A8* are down-regulated while macrophage related genes such as *APOE*, *FBP1* and *FTL* are up-regulated (**Figure 3D**), suggesting the recruitment of infiltrating monocytes and subsequent macrophage differentiation.

**Figure 3.**
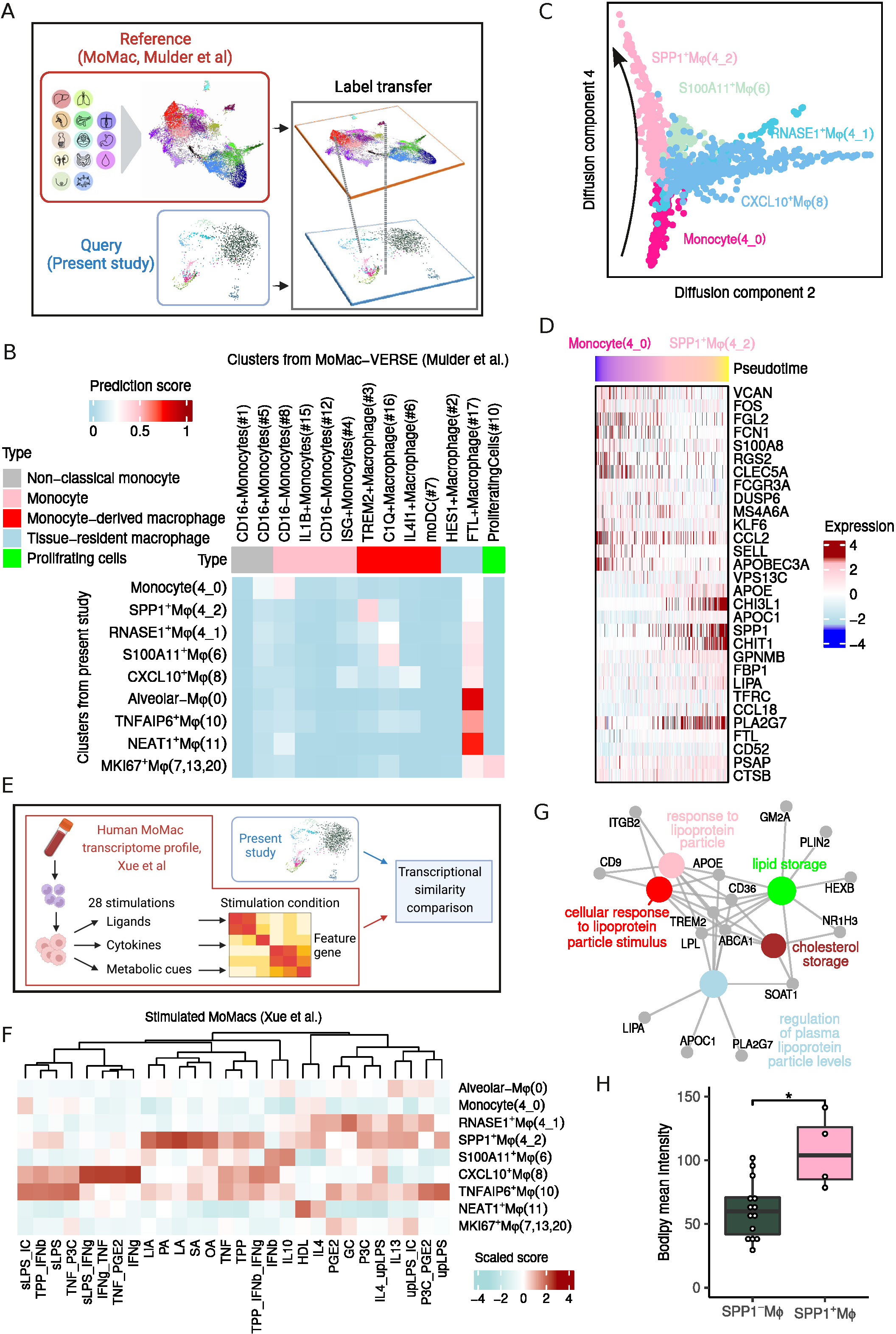
Monocyte derived SPP1^+^Mφ were characterized by up-regulated metabolic pathway. (A) Schematic workflow of transferring the macrophage labels from MoMac dataset to macrophages from present study. (B) Heatmap of transcriptional similarity between macrophages determined in present study and macrophages from the MoMac dataset. (C) Diffusion map embedding of monocyte and monocyte derived macrophages. (D) Differentially expressed genes along pseudotime from monocyte to SPP1^+^Mφ. (E) Schematic workflow of comparing the transcriptional similarity between macrophages from present study and activated macrophages from Xue et al. (F) Heatmap of transcriptional similarity between macrophages determined in present study and activated macrophages from Xue et al. (G) Enriched pathways in SPP1^+^Mφ. (H) Comparison of bodipy intensity in SPP1^-^Mφ and SPP1^+^Mφ. \**p* < 0.05, t-test.

As SPP1^+^Mφ are enriched during early-IPF, we next sought to explore which extracellular cues would specify monocyte to SPP1^+^Mφ development. We made use of a previously published resource that profiled the transcriptomes of human monocyte-derived macrophages under 28 different stimuli *in vitro* (**Figure 3E)** (Xue et al., 2014). Comparison of the transcriptomic similarity between our BALF macrophage subpopulations and this macrophage polarisation spectrum revealed that CXCL10^+^Mφ and S100A11^+^Mφ shared similar profiles to macrophages stimulated by IFN-γ. Notably, SPP1^+^Mφ were predicted to be similar to macrophages stimulated with fatty acids such as lauric acid (LA), oleic acid (OA), linoleic acid (LiA), palmitic acid (PA) and stearic acid (SA) (**Figure 3F**). Consistent with their predicted polarisation via exposure to fatty acids, gene enrichment analysis revealed upregulated lipid localization and lipid storage pathways in SPP1^+^Mφ (**Figure 3G, Figure S4B, Table S5**). Furthermore, BODIPY staining analysis of BALF cells also showed the significant accumulation of lipid droplets in the SPP1^+^Mφ comparing to the SPP1^-^Mφ (**Figure 3H, Figure S4C**).

Together, these data suggest that monocytes are exposed to local lipid-mediated signals in the lung or bronchioalveolar space during differentiation, resulting in the acquisition of the SPP1^+^Mφ transcriptional state, which is associated with lipid metabolism.

### The significant enrichment of SPP1 **^+^**M**φ** in the early-IPF was validated in an independent cohort

Previous studies characterizing lung biopsy samples have reported a profibrotic macrophage population enriched in the end stage of IPF compared to controls (Adams et al., 2020; Morse et al., 2019). Accordingly, we sought to assess whether SPP1^+^Mφ identified in the bronchoalveolar space are analogous to those profibrotic macrophages previously reported. We integrated macrophages from our study along with macrophages from three published IPF datasets (**Figure S5A-B**) (Adams et al., 2020; Habermann et al., 2020; Morse et al., 2019). This joint embedding of 260,747 macrophages revealed the similarity between SPP1^+^Mφ identified in present study and the cluster 3 from the integration analysis, of which was previously determined as SPP1^hi^ macrophage or IPF-expanded macrophage (IPFeMφ) (**Figure S5C-D**). In line with previous studies, quantification of macrophages from lung samples revealed significant enrichment of SPP1^+^Mφ in IPF comparing to control (**Figure S5E**).

Given the strong association of SPP1^+^Mφ and IPF, we sought to quantify the SPP1^+^Mφ in bronchoalveolar lavage liquid samples using flow cytometry analysis, a routine technique used to characterize BALF samples in clinical practice. To determine FACs surface markers for the SPP1^+^Mφ we made use of a recently published CITE-seq data that profiled whole transcriptome and 277 antibodies of BAL cells from COVID-positive patients as well as control samples (**Figure 4A**) (Bosteels et al., 2022). Integration of our macrophage subpopulations to the CITE-seq has revealed that RNASE1^+^Mφ and SPP1^+^Mφ showed strong similarity to cluster 1 and cluster 10 from the CITE-seq dataset, respectively (**Figure S5F-H**). These two populations could be potentially separated from alveolar macrophages by antibody CD10 (**Figure 4B**). In addition, DEG analysis has revealed that FOLR2 and ALCAM (CD166) is uniquely highly expressed in RNASE1^+^Mφ and SPP1^+^Mφ, respectively (**Figure 4C, Figure S5G**). Therefore, we designed a modified spectral flow cytometry panel to characterize macrophages subsets within BALF samples. This panel, for the first time, allowed for the effective isolation of true alveolar macrophages (CD169^+^CD206^+^CD10^+^FOLR2^-^), RNASE1^+^Mφ (interstitial macrophages, CD169^+^CD206^+^CD10^-^ FOLR2^+^) and SPP1^+^Mφ (CD169^+^CD206^+^CD10^-^FOLR2^-^CD166^+/-^) within human BALF (**Figure 4D**).

**Figure 4.**
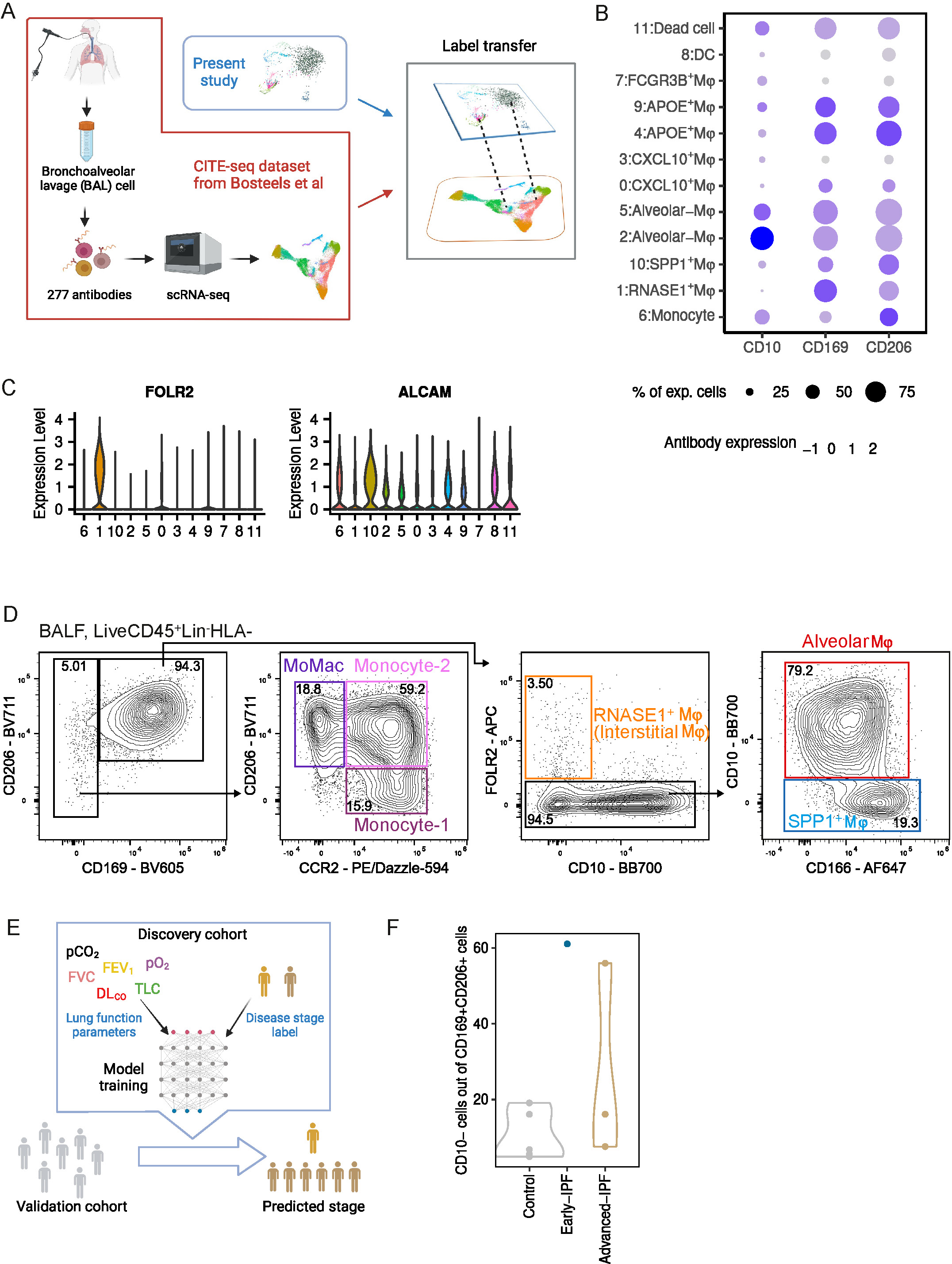
Enrichment of SPP1^+^Mφ in the early-IPF group was validated in an independent IPF cohort. (A) Schematic workflow of integrating macrophages from present study to the CITE-seq dataset to determine antibodies for SPP1^+^Mφ. (B) Antibody (CD10, CD169 and CD206) expression level in all clusters from the CITE-seq dataset. (C) Gene (ALCAM and FOLR2) expression level in all clusters from the CITE-seq dataset. (D) Strategy to gate macrophage subpopulations in BALF samples for the Spectrum CD10 + Extended mac panel. (E) Schematic workflow of machine learning based analysis to stratify IPF patients from the validation cohort into early- and advanced-IPF groups. (F) Frequency of CD10-populations in the validation cohort separated by disease stage groups. \**p* < 0.05, \*\**p* < 0.01, *** *p* < 0.001, paired t-test.

With this newly developed panel we profiled a validation cohort which contains 7 IPF patients and 7 control samples. We first applied machine-learning based analysis to predict disease stage of IPF patients in the validation cohort, by using patients listed in the Figure 1 as the training data, termed as discovery cohort (**Figure 4E**). This analysis has determined 1 early-IPF and 6 advanced-IPF patient. Embedding patients from both discovery and validation cohort has further supported that the early-IPF patient is at early stage of disease development (**Figure S6A**). Despite of the low number of patients in the early-IPF, quantification of CD10^-^macrophage populations we indeed observed a promising increased trend of SPP1^+^Mφ at the early stage of disease development (**Figure 4F, Table S6**).

In summary, these data have shown that the SPP1^+^Mφ identified in the bronchoalveolar space in our study showed similar transcriptional profile as one profibrotic macrophage population previously reported in the lung tissue samples from patients with end stage of IPF. Moreover, we were able to quantify this population in BALF samples using a novel FACs panel, providing an interesting possibility to detect SPP1^+^Mφ in clinical practice.

## Discussion

IPF is a rare fibrotic lung disease with unfavourable treatment outcomes. Previous studies profiled the end stage of transplanted lung samples have demonstrated cellular markers related to IPF (Adams et al., 2020; Ayaub et al., 2021; Habermann et al., 2020; Morse et al., 2019). However, due to the significant morbidity and mortality caused by surgical lung biopsy procedures, these markers were difficult to be applied in the clinical practice (Hutchinson et al., 2015). Hence, cellular markers from easy to access samples to determine IPF at the early stage are urgently demanded.

Here, we used unsupervised hierarchical clustering to determine patients at early stage of the disease, of which showed similar lung function as control patients but with fibrosis in the lung. Single-cell whole transcriptome analysis of the BALF samples has demonstrated the great macrophage heterogeneity in the IPF bronchoalveolar space. Quantification of macrophage subpopulations has revealed that SPP1^+^Mφ were prominently enriched at the early stage of IPF. Functional analysis *in silico* has suggested that SPP1^+^Mφ were derived from monocytes upon fatty acid stimulation. Additionally, we have established a FACs panel to quantify SPP1^+^Mφ in BALF samples, of which was used to profile a validation cohort and confirmed the pronounced accumulation of SPP1^+^Mφ at the early stage of IPF. Together these data have shown that SPP1^+^Mφ could be potentially used as a cellular marker to determine patients at the early stage of IPF in clinical practice.

SPP1^+^Mφ were featured by gene *TREM2* and were previously termed as TREM2^+^Mφ, accumulation of which occurs at early/mild stage of development in other diseases. For instance, by mapping the macrophages from BALF samples from COVID patients into the MoMac-VERSE, Mulder and colleagues have demonstrated that TREM2^+^Mφ are enriched in patients with mild COVID symptoms (Liao et al., 2020; Mulder et al., 2021). Additionally, a recent work on murine heart failure model has suggested that the frequency of TREM2^hi^SPP1^hi^Mφ was gradually increased after the induction of myocardial infarction and was peaked at day 5, but was decreased at day 7 (Jung et al., 2022). In line with this work, here we have shown that SPP1^+^Mφ were found at low levels in the control patients but were significantly enriched in the early-IPF group. These data suggested that SPP1^+^Mφ might play an important role in responding to, or driving, the pathological factors at early stage of the disease progression.

Analogues of SPP1^+^Mφ have been determined in multiple human and murine diseases and tumors, however the impact of SPP1^+^Mφ on disease and tumor response is context specific. For example, in the murine carcinoma and ovarian cancer, TREM2^+^Mφ are associated with increased tumor growth and immunosuppressive response (Binnewies et al., 2021; Katzenelenbogen et al., 2020; Molgora et al., 2020; Park et al., 2023). In contrast, recent work in hepatocellular carcinoma and skin transplantation demonstrated that TREM2^+^Mφ might limit tissue fibrosis and promote wound healing (Esparza-Baquer et al., 2021; Henn et al., 2023). The pathological function of SPP1^+^Mφ in human IPF remains largely unknown. In our study we have suggested that SPP1^+^Mφ were derived from monocytes upon fatty acid stimulation. This result is in agreement with previous proteomic analysis of BALF samples showing that IPF bronchoalveolar space is enriched with palmitic acid and stearic acid fatty acids (Chu et al., 2019; Mayr et al., 2020). Therefore future studies to investigate how the microenvironment at early state of IPF induced the accumulation of the SPP1^+^Mφ would help to understand the pathogenic mechanisms of IPF progression.

One limitation in our study is that the low number of IPF patients might impair the statistical comparison of SPP1^+^Mφ among the groups. However, by integrating datasets from three published studies we were able to confirm that SPP1^+^Mφ were indeed significantly increased in IPF samples, although quantification of SPP1^+^Mφ in early-IPF is not possible due to the lack of early-IPF samples in the literature. Future collaborative studies from multiple centres could offer the opportunity to systematically investigate the prognostic value of SPP1^+^Mφ.

Collectively in this study we have shown that monocyte derived SPP1^+^Mφ were predominantly enriched in the early stage of IPF. We have also provided a FACS panel to quantify SPP1^+^Mφ in BALF samples, which could be applied in future clinical practice to facilitate early diagnosis, and aid patient prognosis.

## Resource Availability

### Lead Contact

Further information and requests for resources and reagents should be directed to and will be fulfilled by the Lead Contact, Andreas Schlitzer.

### Material Availability

This study did not generate new unique reagents.

### Data and code availability

scRNA-seq data generated during this study are deposited at the National Center for Biotechnology Information (NCBI) under the number GEO xxx. R scripts used for the analysis have been deposited on GitHub (https://github.com/jiangyanyu/IPF_2023). Scripts were executed in the docker image: https://hub.docker.com/r/jiangyanyu/jyu_r4.1.2. Seurat object for the analysis has been deposited on GEO xxx.

### Experimental Model and Subject Details

This study was approved by the Institutional Review board of the University Hospital Bonn. All individuals provided written informed consent according to the Declaration of Helsinki before samples were collected.

## Method Details

### Isolation of cells from bronchoalveolar lavage fluid (BALF)

Human BALF samples were obtained from University Hospital of Bonn and Marien Hospital Bonn. All samples were processed within two hours after collection and all steps were performed at 4°C. The fluid was filtered through a 100 μM cell strainer (Merck) and was centrifuged at 350 g for 7 min. The supernatant was immediately frozen in liquid nitrogen and was later stored in -80 °C. BALF cell pellets were washed once with flow cytometry staining buffer (FACS buffer, PBS supplemented with 2 mM EDTA and 0.5% fetal calf serum). Erythrocytes were removed using the red blood cells lysis buffer (Biolegend). Cells were washed once and resuspended in FACS buffer for further analysis.

### Flow cytometry analysis of BALF samples from the discovery cohort

BALF cells were isolated and prepared as described above, then were incubated with fixable viability dye (1:1000, BioLegend) for 15 min at room temperature in the dark. After a washing step, cells were stained with the antibody cocktail for one hour on ice in the dark (Table S7). Later cells were washed once and were incubated with streptavidin-PE (1:200, BioLegend) for 20 min on ice in the dark. After another washing step, cells were resuspended in FACS buffer and were acquired using the FACSymphony Cell Analyzer (BD Biosciences). Data were analyzed using FlowJo software (v10.8.1, BD Biosciences).

### Flow cytometry analysis of BALF samples from the validation cohort

BALF cells were isolated and prepared as described above. After one washing step cells were resuspended in the blocking buffer and were incubated at 37°C for 15 min. Then cells were stained with CCR7 (3:100, BioLegend) and CADM1 (2:100, BioLegend) at 37°C for 25 min. After one step washing with cold PBS, cells were incubated with fixable viability dye (1:1000, BioLegend) for 20 min at room temperature in the dark. Later cells were washed with cold PBS and were incubated with antibody cocktail for 30 min on ice in the dark (Table S8). After another washing step cells were resuspended in cytofix buffer (BioLegend) and were incubated on ice for 20min. Finally after removing the cytofix buffer, cells were resuspended in FACS buffer and were acquired using the SA3800 Spectral Analyzer (SONY). Data were analyzed using FlowJo software (v10.8.1, BD Biosciences).

### Seq-Wel based scRNA-seq

Seq-Well based scRNA-seq was performed as previously reported (Hughes et al., 2020). The Seq-Well arrays were generated and functionalized in the PRECISE platform for single cell genomics and epigenetics in the Deutsches Zentrum für Neurodegenerative Erkrankungen (DZNE) according to steps previously described (Bassler et al., Frontiers in Immunology, 2022). After loading 110,000 mRNA capture beads (Chemgenes Corp) to the functionalized array, 20,000 BALF cells were loaded into the array. Once cells were settled into wells, the array was washed five times with cold PBS and was subsequently sealed with hydroxylated polycarbonate membrane (Sterlitech) with a pore size of 10 nm. The sealed array with loaded cells was incubated at 37°C for 30 min and was later placed in lysis buffer (5 M guanidine thiocyanate, 1 mM EDTA, 0.5% Sarkosyl and 1% β-mercaptoethanol in H_2_O) for 20 min at room temperature. After cell lysis the array was placed in hybridization buffer (2 M NaCl, 3 mM MgCl_2_ and 0.5% Tween-20 in PBS) at 4°C for 40 min. The mRNA capture beads were collected by contrifudging at 2,000 g for 5 min, and was immediately subjected for reverse transcription reaction (Maxima RT buffer, 12% PEG8K, 1 mM dNTPs, 1 U/µL RNAse inhibitor, 2.5 µM template switch oligo (TSO) and 10 U/µL Maxima H-RT enzyme in H_2_O) for 30 min at room temperature and 90 min at 52°C with end-over-end rotation. The obtained bead-bound cDNA product was washed once with TE-TW (0.1% Tween-20, 1 mM EDTA and 10 mM Tris-HCl pH 8.0 in H_2_O), once with TE-SDS (0.5% SDS, 1 mM EDTA and 10 mM Tris-HCl pH 8.0 in H_2_O) and twice with TE-TW, and was resuspended in TE-TW solution for following steps.

A second-strand synthesis was performed before cDNA amplification and sequencing. First, to remove excess primers the bead-bound cDNA was treated with exonuclease I (Exonuclease buffer and 1 U/µL Exonuclease in H_2_O) for 50 min at 37°C. After washing once with TE-SDS and twice with TE-TW, the beads were solvated in 0.1 M NaOH for 5 min at room temperature with end-over-end rotation. Following another sequential wash once with TE-TW and once with TE buffer (1 mM EDTA and 10 mM Tris-HCl pH 8.0 in H_2_O), beads were subjected for the second-strand synthesis reaction (Maxima RT buffer, 12% PEG8K, 1 mM dNTPs, 2 uM dN-Smart randomer and 1 U/µL Klenow Fragment in H_2_O) for 1 hour at 37°C with over-end-over rotation. Next the beads were washed twice with TE-TW and once with TE, and were resuspended in water for following steps.

After second-strand synthesis the beads underwent whole transcriptome amplification (WTA) reaction. Briefly, beads were first counted with Fuchs-Rosenthal cytometer (Merck). Parallel PCR reactions were performed to make sure to amplify all beads from the entire array. A portion of 5,000 beads were subjected for each reaction (KAPA HiFi Hotstart readmix and SMART primer in H_2_O) using cycling conditions: 95°C for 3 min, then 4 cycles of 98°C for 20 seconds, 65°C for 45 seconds, and 72°C for 3 min, followed by 12 cycles of 98°C or 20 seconds, 67°C or 20 second, and 72°C for 3 min, and lastly a final extension of 72°C for 5 min. The PCR products were combined into small pools, with each pool containing 10,000-12,000 beads. Usually an array generated 3 to 4 WTA pools, which were subjected to following purification and sequencing steps in parralel. Next the pooled products underwent two sequential purification steps using AMPure XP beads (Beckman Coulter) with 0.6x and 0.8x volumetric ratio according to the manufacture’s instruction. The quality of purified WTA products was assessed using Tapestation 2200 (Agilent) with high sensitivity D5000 assay, and WTA products were quantified using Qubit 4 Fluorometer (Thermo Fisher Scientific) with high-sensitivity dsDNA assay.

Next purified WTA products were subjected for library construction and sequencing. In brief, 200 pg of purified WTA products from each pool were subjected for library construction with custom designed New-P5-SMART-PCR primer using the Nextera XT DNA library preparation kit (Illumina) following the detailed protocol provided by the manufacture. After purification and quantification the sequencing library was diluted to 1.25 nM for sequencer loading. Sequencing was performed in paired-end mode (21 cycles for read 1 using custom designed read 1 primer, 8 cycles for i7 index and 61 cycles for read 2) with Novaseq S2 flow cells (100 cycles) using Novaseq 6000 sequencer (Illumina) in the Life and Brain Center, Bonn.

### Immunofluorescence assay to quantify cholesterol levels

After isolated from BALF samples as mentioned above, 100, 000 BALF cells were resuspended in 1.5 mL RPMI medium (PNA-Biotech), and were seeded on the coverslip that was placed in one well of the 6-cell culture dish. After seeding, cells were incubated at 37°C for 2 hours and were washed once with PBS. Then cells were fixed with 4% PFA (Thermo Fisher Scientific) for 20 min at room temperature and were transferred to a humid chamber. After 3 times washing steps with PBS, cells were incubated in the blocking buffer (3% BSA in PBS) for 1 hour at room temperature. Later cells were stained with SPP1 (1:200, Sigma-Aldrich) and CD206 (1:100, Bio-Rad) at 4°C overnight. After 3 steps of washing with PBS and one step washing with HBSS, cells were stained with BODIPY-cholesterol (2.5 µM in HBSS) and secondary antibodies at 4°C for one hour. After 3 steps of washing with PBS, cells were stained with DAPI (1:2000, BioLegend) for 20 min at room temperature and later were washed 3 times with PBS. After removing residual solutions on the coverslip, samples were mounted with FluoromountG (Thermo Fisher Scientific) medium. Cells were imaged using 405 nm, 488 nm, 561 nm and 640 nm laser lines from the Zeiss LSM 889 Airyscan system (Zeiss). Fluorescent intensities of each marker for each BALF cell were quantified using Fiji (v2.9.0).

## Quantification and Statistical Analysis

### Unsupervised hierarchical clustering of the discovery cohort

Pulmonary function test parameters were used to perform the cluster analysis. After removing the highly correlated parameters, six pulmonary function test parameters were selected for the following cluster analysis. Missing values were imputed by R package Amelia (v1.8.1). The imputed matrix was subjected to compute Euclidean distances by dist from R package stats (v4.1.3). Simultaneously, the distance matrix was used to generate a cluster dendrogram using hclust from R package stats (v4.1.3).

### Pre-processing of scRNA-seq data

Demultiplexed fastq files from sequencer were loaded to a snakemake-based data pre-processing pipeline (v0.31) which was provided by the McCarrol lab (Macosko et al., 2015). Reads were aligned to human reference genome (Ensembl build 38 release 91). Count matrices generated from the pipeline were imported into R (v4.1.3) for further analyses. Before data integrating cells with low quality (UMIs per cell < 700 or mitochondrial gene percentage > 40% or genes per cell < 500) were removed according to the filtering criteria previously reported in the Seq-Well paper (Hughes et al., 2020).

### Integration of cells from present study and cells from Bassler et al

Reciprocal PCA based approach wrapped in Seurat package (Hao et al., 2021) was used to integrate cells from present study and BALF cells from six healthy individuals generated by Seq-Well pipeline from a recently published study (Baßler et al., 2022). For dataset from each patient or control individual, normalization and feature genes (top 2,000) determination were performed independently using default settings of NormalizaData and FindVariableFeatures, respectively. Then independent scaling and PCA analysis of each dataset were performed using features that were repeatedly variable across all datasets defined by SelectIntegrationFeatures. After identifying anchors across all datasets using FindIntegrationAnchots by setting reduction as rpca, integration was performed by IntegrateData.

Downstream dimension reduction and clustering were performed on the corrected data stored in the integrated assay. In short, after scaling and PCA analysis of corrected data, first 30 PCs were used to construct a shared nearest-neighbor graph which was loaded for clustering that was performed with resolution of 1 to generate 26 clusters. Two clusters (cluster 1 and 18) featured only by mitochondrial genes were removed from downstream analysis. Doublet cells (cluster 2, 3, 23 and 25) were identified using R package scDblFinder (v1.8.0) and were removed from downstream analysis. The UMAP based dimension reduction was performed to visualize cells and clusters in a low dimension, by using the first 30 PCs. To gain a better overview of macrophage subpopulations, all macrophage clusters were selected for UMAP analysis using RunUMAP with the embedding parameter b as 0.9.

### Cell type annotation comparison between present study and human lung atlas

Cell types annotated by canonical markers were compared to annotations from the human lung atlas (Travaglini et al., 2020). In brief, we first downloaded the list (Table S4 from the Travaglini et al. study) containing marker genes of all clusters. Marker genes (avg_logFC > 1) derived from 10x Chromium (sheets name without “SS2”) were subjected to AddModuleScore function from Seurat package, to calculate average expression (prediction score) of these marker genes in each cell from present study. The average prediction score was scaled and visualized in heatmap using ComplexHeatmap package (v2.10.0) (Gu et al., 2016).

### Differentially expressed genes identification

Differentially expressed genes (DEGs) between clusters were determined by performing a Wilcox test on normalized data before integration using FindAllMarkers function in Seurat. DEGs were defined as avg_log2FC > 0.5 and p_val_adj < 0.001.

### GO enrichment analysis

To identify whether cluster is enriched with genes related to certain biological pathways, GO enrichment analysis was performed using the R package clusterProfiler (v.4.2.2). In brief, DEGs for each cluster were submitted to enrichGo function. All genes present in the count matrix were served as background genes. GO terms with p-adjust < 0.001 were considered as enriched in the cluster.

### Label transferring between macrophages from present study and cells from MoMac-VERSE

To transfer the labels from the MoMac-VERSE database to macrophages from present study, TransferData function from Seurat package was used. In brief, macrophages from lung samples in the MoMac-VERSE were isolated and were used as the reference. Then FindTransferAnchors function was used to find anchors between the reference and query dataset (macrophages from present study) using the first 30 PCs. After finding anchors, TransferData function was used to classify the query cells based on the reference data. Prediction scores from last step was added to the query metadata. The average prediction score per cluster was visualized in heatmap using ComplexHeatmap package (v2.10.0).

### Transcriptional similarity comparison between macrophages from present study and macrophages from human activated macrophage spectrum

To determine the condition that would differentiate monocyte into the SPP1^+^Mφ, we compared the transcriptional similarity between macrophages from current study to activated macrophages from Xue et al. In brief, we first downloaded the list containing marker genes for each activated macrophage population from the paper. Marker genes for each activate macrophage population were subjected to AddModuleScore function from Seurat package, to calculate average expression (prediction score) of these marker genes in each cell from present study. The average prediction score was scaled and visualized in heatmap using ComplexHeatmap package (v2.12.1).

### Integration of macrophages from present study and macrophages from four previous studies

To determine whether SPP1^+^Mφ present in lung biopsy samples, we integrated macrophages from present study to macrophages from three previously published studies (Adams et al., Haberman et al., Morse et al.). First, we selected macrophages from IPF patients and control samples from Adams et al. (“Manuscript_Identify” as “Macrophage” or “Macrophage_Alveolar”), Haberman et al. (“celltype” as “Macrophages” or “Proliferating Macrophages”) and Morse et al. (clusters expressing MARCO, CD68 and FABP4). The integration process was performed using Seurat package. In short, for each of the four datasets, normalization and feature genes (top 2,000) determination were performed independently using default settings of NormalizaData and FindVariableFeatures, respectively. Then independent scaling and PCA analysis of each dataset were performed using features that were variable across all datasets defined by SelectIntegrationFeatures. After identifying anchors across all datasets using FindIntegrationAnchors by setting reduction as rpca, integration was performed by IntegrateData. Downstream dimension reduction and clustering were performed on the corrected data stored in the integrated assay. In short, after scaling and PCA analysis of corrected data, first 30 PCs were used to construct a shared nearest-neighbor graph which was loaded for clustering that was performed with resolution of 0.3 to generate 18 clusters.

To determine the transcriptional similarity between SPP1^+^Mφ and previously reported profibrotic macrophage, we downloaded the feature genes of these macrophage populations (sheet 3 of Table S3 from Wendisch et al., Cell, 2021). Then the list of feature genes was subjected to the AddModuleScore function from Seurat package, to calculate average expression (prediction score) of these marker genes in each cell from the integrated dataset.

### Reanalysis of CITE-seq dataset

The CITE-seq data was downloaded from the link provided by the recently published study (Bosttels et al., Cell reports medicine, 2022). First, we selected mononuclear phagocytes from the CITE-seq dataset (“Annotation” as “cDC”, “Macrophage” or “Macrophage:Alveolar”). Analysis of the CITE-seq data was performed by using Seurat package. In brief, we first generated a Seurat object containing the gene expression matrix and antibody expression matrix for each patient from the CITE-seq dataset. Then the list of objects was subjected to RunFastMNN function from the SeuratWrappers package to remove batch effects. Later, first 30 PCs were used to construct a shared nearest-neighbor graph which was loaded for clustering that was performed with resolution of 0.6 to generate 11 clusters.

To identify the SPP1^+^Mφ in the CITE-seq dataset, we transferred the labels of macrophages from the present study (reference) to the CITE-seq dataset (query) by using FindTransferAnchors and TransferData functions from Seurat package, as described above. Prediction score for each cell was added to the query metadata. The average prediction score per cluster was visualized in heatmap using ComplexHeatmap package (v2.12.1).

To determine surface markers for SPP1^+^Mφ for FACs analysis, we used FindMarkers function from Seurat package. Cells in the cluster 10 were set as ident.1, while rest of the cells were set as ident.2. Differentially expressed antibodies were visualized in a volcano plot.

### Trajectory analysis

To infer the trajectory from monocytes to macrophages, monocytes and monocyte derived macrophages from present study were embedded on a diffusion map using the Destiny package (v3.14.0). Briefly, the first 50 PCs were subjected to find_sigmas function to find optimal sigma. Then DiffusionMap function was used to compute diffusion components and corresponding eigenvectors with k set as 5. Cells were embedded on 2D space using diffusion component 2 and 4.

### Machine-learning based analysis to stratify the validation cohort

To stratify the IPF patients from the validation cohort, machine learning based analysis was performed using caret package (v6.0.94). In short, 75% of the IPF patients from the discovery cohort was used to train the model and the rest of 25% was used to test the accuracy of prediction. Decision tree (rpart) based method was used for the model training. Lastly, the validation cohort was subjected to predict function of the caret package.

### Quantification and statistical analysis

Statistical analysis of flow cytometry data was performed using Prism 9 (Graphpad) and R (v4.1.3). respectively. If not otherwise stated, the statistical analysis was performed according to total sample size. A t-test was used if samples are less than 10, otherwise a Wilcoxon rank-sum test was used. All workflow figures were created with Biorender.com.

## Acknowledgments

We thank all patients and families involved in this study. We thank Jonas Schulte-Schrepping, Kristian Händler, André Heimbach, Collins Osei-Sarpong and Heidi Theis for sequencing assistance. We thank Marie Vandestienne and Wenming Huang for critical feedback on the manuscript. This work was supported by the German Research Foundation to D.S., J.Y. and A.S.; EXC ImmunoSensation2 to J.Y., J.T., J.H. and A.S.; Emmy Noether Research Grant to A.S.; EXC Hausdorff Center for Mathematics to J.H.; the SFB 1454 for A.S., and J.H; and the University of Bonn via the Schlegel professorship to J.H.

## Author contributions

Conceptualization, J.Y. and A.S.; methodology, J.Y., J.T., J.H., M.Z., and L.Z.; data analysis, J.Y., J.T., and M.B.; resources, C.P., L.B., T.S., W.S., J.S., J.H. and D.S.; writing, J.Y. and A.S.

---

**Figure S1.**
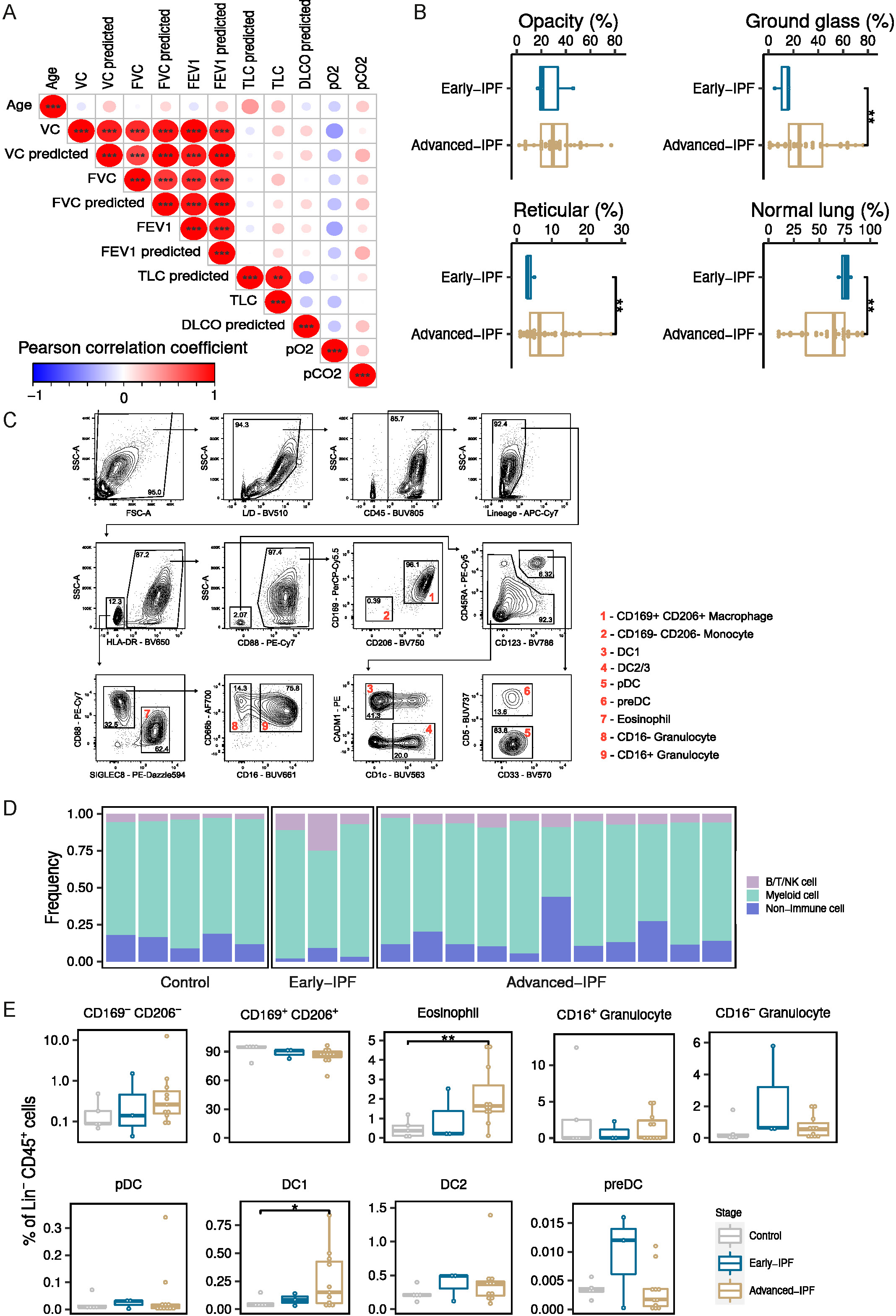
Unsupervised hierarchical clustering stratified IPF patients into two groups. (A) Pearson’s correlation coefficient of lung function parameters. (B) Boxplot of lung function parameters from high-resolution computed tomography (HRCT) images between early- and advanced-IPF groups. \*\**p* < 0.01, paired t-test. (C) Strategy to gate immune cells in BALF samples. (D) Composition of cells in BALF samples determined by flow cytometry analysis. (E) Composition of myeloid cells in all samples grouped by stages.

**Figure S2.**
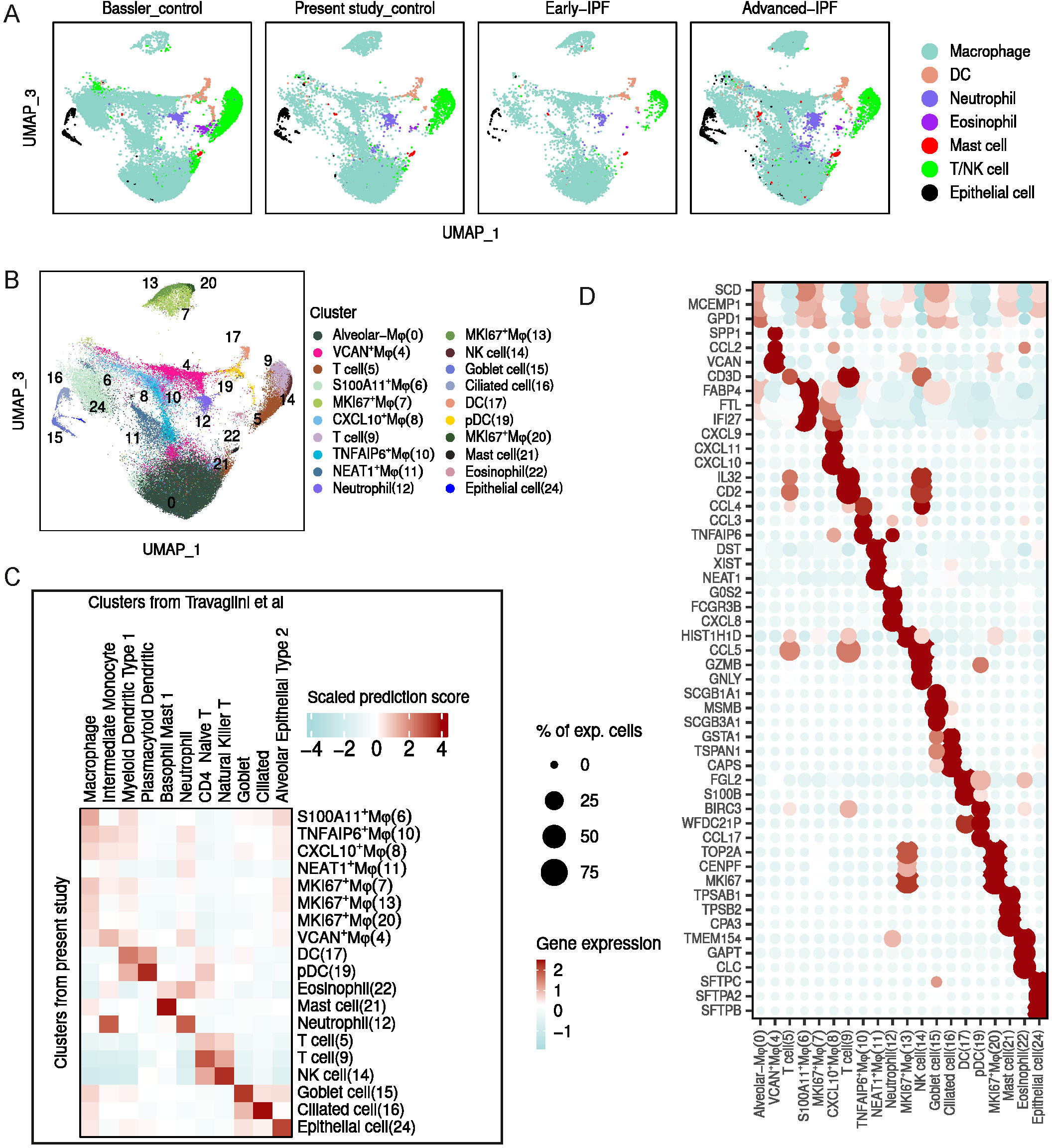
Single cell transcriptomic analysis revealed macrophage heterogeneity in the bronchoalveolar space. (A) UMAP embedding of cells from BALF samples grouped by source of control samples and disease groups. (B) UMAP embedding of cell populations determined in BALF samples. (C) Transcriptional similarity between cell types determined in present study and human lung cell atlas. (D) Top 5 feature genes of each cluster. Feature genes threshold: avg_log2FC >1, p_val_adj <0.001.

**Figure S3.**
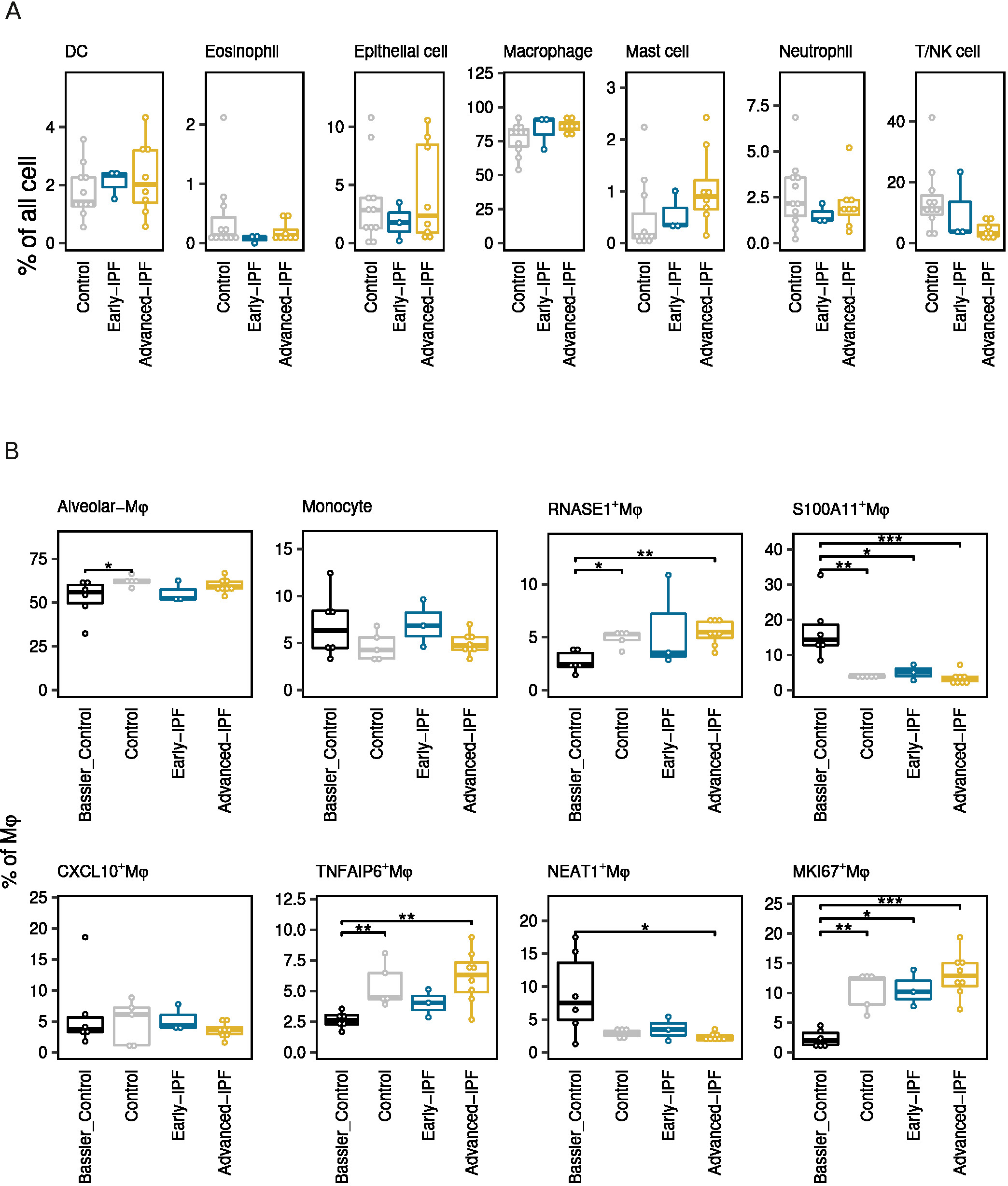
SPP1^+^Mφ were predominantly accumulated at the early stage of IPF development. (A) Frequency of major cell population in the control and disease groups. (B) Frequency of each subpopulation in the control and disease groups. \**p* < 0.05, \*\**p* < 0.01, *** *p* < 0.001, Wilcox-test.

**Figure S4.**
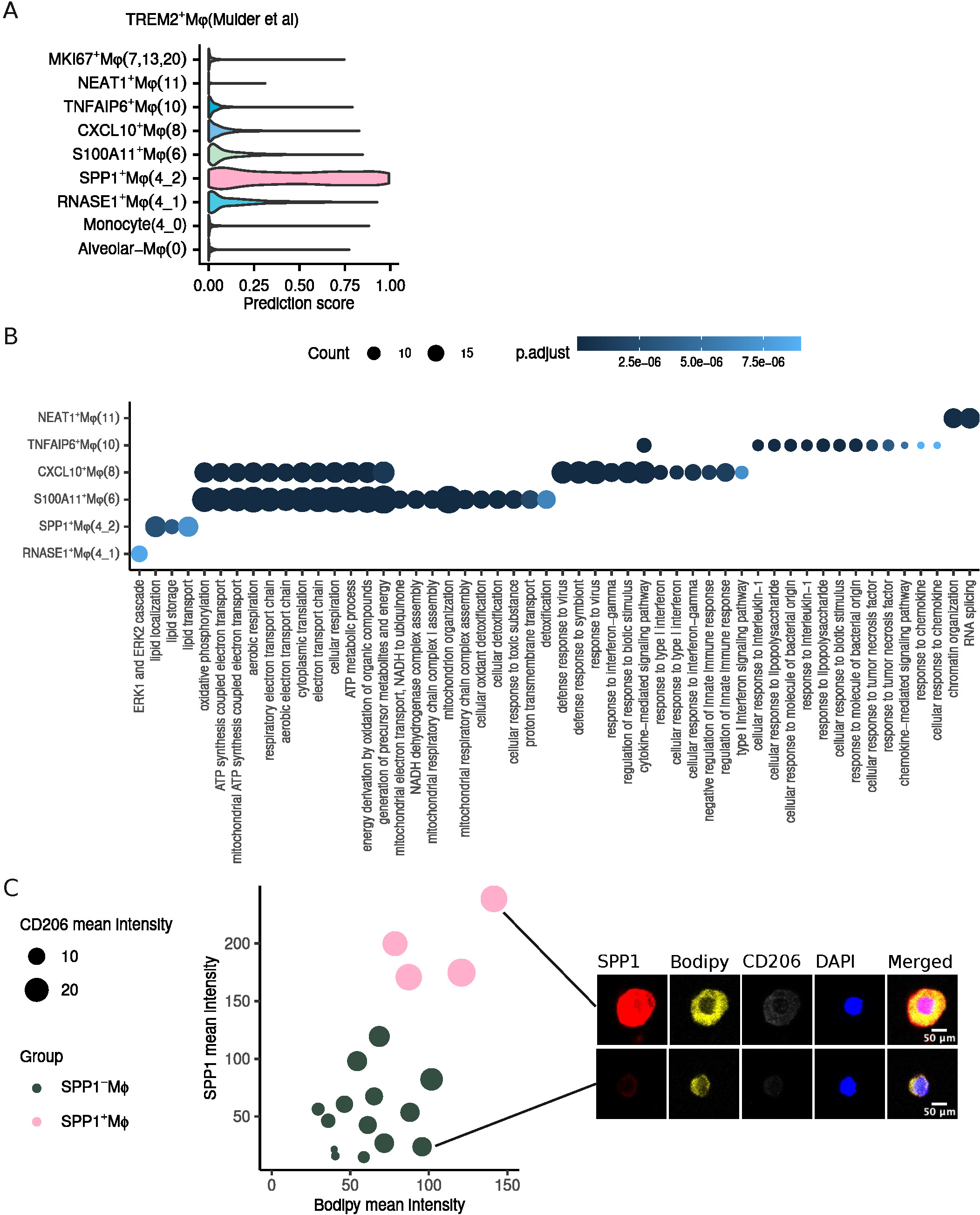
Monocyte derived SPP1^+^Mφ were characterized by up-regulated metabolic pathway. (A) Transcriptional similarity between macrophage subpopulations in current study and TREM2^+^Mφ from the MoMac database. (B) Enriched GO terms in macrophage subpopulations. (C) Quantification of bopidy intensity in SPP1^-^Mφ and SPP1^+^Mφ. Examples of SPP1^-^Mφ and SPP1^+^Mφ are shown on the right.

**Figure S5.**
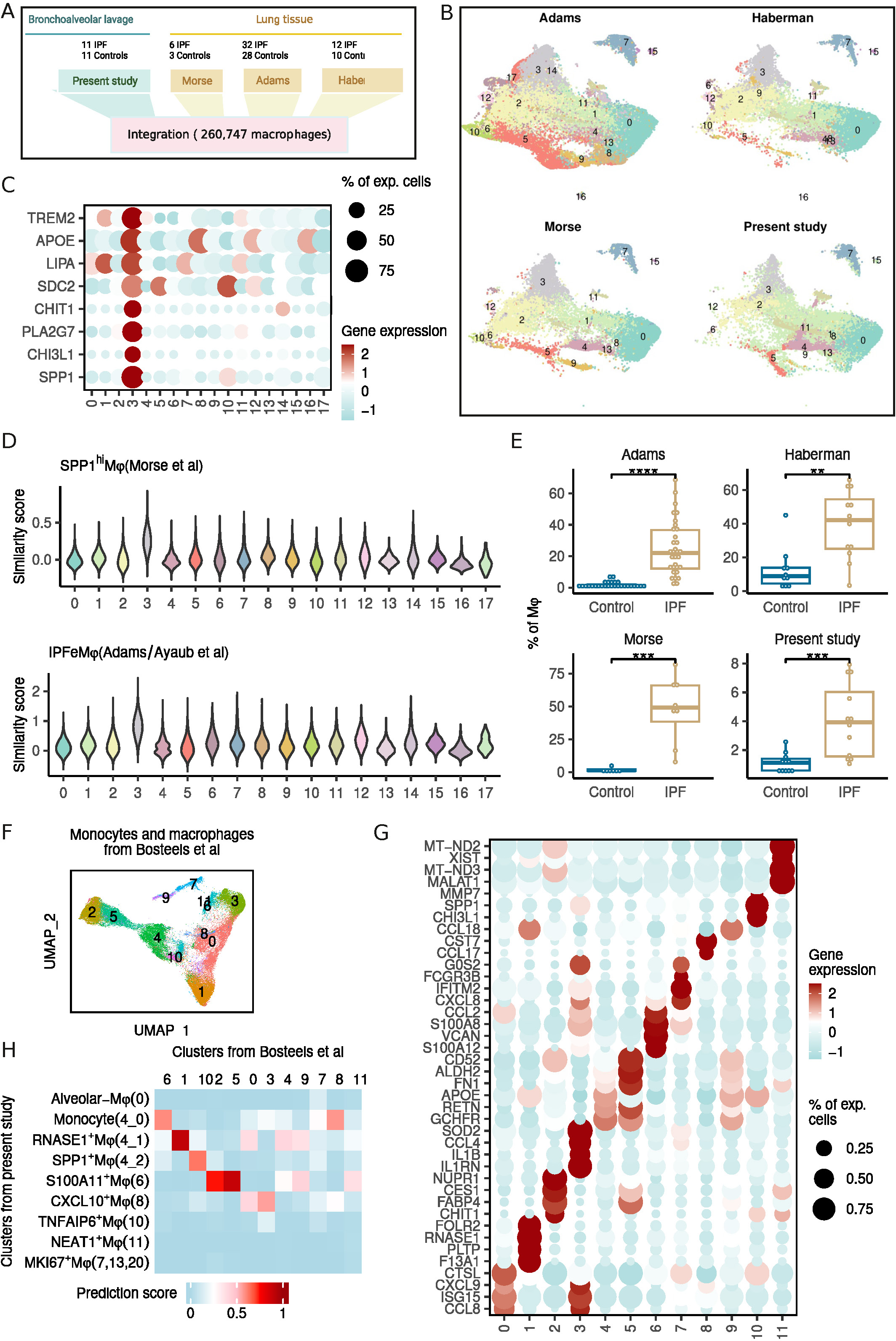
SPP1^+^Mφ were significantly enriched in lung biopsy samples from patients with end-stage of IPF. (A) Schematic workflow of integrating macrophages from present study to macrophages from three other published studies. (B) UMAP embedding of macrophages grouped by studies. (C) Expression of feature genes of SPP^+^Mφ from present study on the clusters from the integrated dataset. (D) Transcriptional similarity of SPP^hi^Mφ and IPFeMφ from previous studies on clusters from the integrated dataset. (E) Frequency of cluster 3 from the integrated dataset in control and IPF samples. (F) UMAP embedding of monocytes and macrophages from Bosteels et al. (G) Differentially expressed genes for clusters from Figure S5F. (H) Heatmap of transcriptional similarity between macrophages determined in present studies and clusters from Figure S5F.

**Figure S6.**
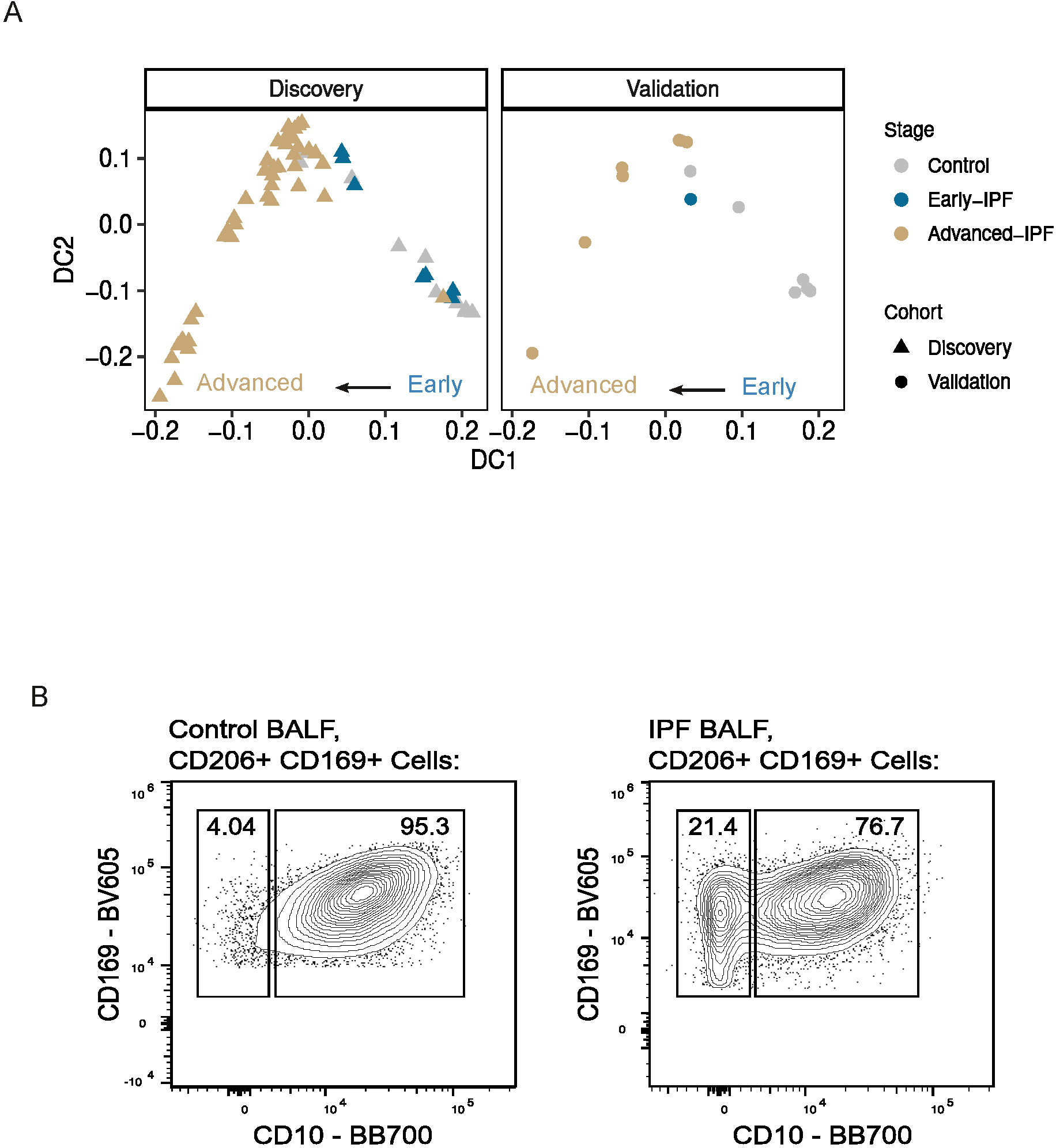
Enrichment of SPP1^+^Mφ in the early-IPF group was validated in an independent IPF cohort. (A) Diffusion map embedding of discovery and validation cohort. (B) Strategy to gate macrophage subpopulations in BALF samples for the Spectrum CD10 panel.

## References

Adams, T.S., Schupp, J.C., Poli, S., Ayaub, E.A., Neumark, N., Ahangari, F., Chu, S.G., Raby, B.A., DeIuliis, G., Januszyk, M., et al. (2020). Single-cell RNA-seq reveals ectopic and aberrant lung-resident cell populations in idiopathic pulmonary fibrosis. Sci. Adv. 6, eaba1983. 10.1126/sciadv.aba1983.

Allden, S.J., Ogger, P.P., Ghai, P., McErlean, P., Hewitt, R., Toshner, R., Walker, S.A., Saunders, P., Kingston, S., Molyneaux, P.L., et al. (2019). The Transferrin Receptor CD71 Delineates Functionally Distinct Airway Macrophage Subsets during Idiopathic Pulmonary Fibrosis. Am. J. Respir. Crit. Care Med. 200, 209–219. 10.1164/rccm.201809-1775OC.

Ayaub, E., Poli, S., Ng, J., Adams, T., Schupp, J., Quesada-Arias, L., Poli, F., Cosme, C., Robertson, M., Martinez-Manzano, J., et al. (2021). Single Cell RNA-seq and Mass Cytometry Reveals a Novel and a Targetable Population of Macrophages in Idiopathic Pulmonary Fibrosis. BioRxiv.

Baßler, K., Fujii, W., Kapellos, T.S., Dudkin, E., Reusch, N., Horne, A., Reiz, B., Luecken, M.D., Osei-Sarpong, C., Warnat-Herresthal, S., et al. (2022). Alveolar macrophages in early stage COPD show functional deviations with properties of impaired immune activation. Front. Immunol. 13.

Binnewies, M., Pollack, J.L., Rudolph, J., Dash, S., Abushawish, M., Lee, T., Jahchan, N.S., Canaday, P., Lu, E., Norng, M., et al. (2021). Targeting TREM2 on tumor-associated macrophages enhances immunotherapy. Cell Rep. 37. 10.1016/j.celrep.2021.109844.

Bosteels, C., Van Damme, K.F.A., De Leeuw, E., Declercq, J., Maes, B., Bosteels, V., Hoste, L., Naesens, L., Debeuf, N., Deckers, J., et al. (2022). Loss of GM-CSF-dependent instruction of alveolar macrophages in COVID-19 provides a rationale for inhaled GM-CSF treatment. Cell Reports Med. 3. 10.1016/j.xcrm.2022.100833.

Carraro, G., Mulay, A., Yao, C., Mizuno, T., Konda, B., Petrov, M., Lafkas, D., Arron, J.R., Hogaboam, C.M., Chen, P., et al. (2020). Single-Cell Reconstruction of Human Basal Cell Diversity in Normal and Idiopathic Pulmonary Fibrosis Lungs. Am. J. Respir. Crit. Care Med. 202, 1540–1550. 10.1164/rccm.201904-0792OC.

Chakarov, S., Lim, H.Y., Tan, L., Lim, S.Y., See, P., Lum, J., Zhang, X.-M., Foo, S., Nakamizo, S., Duan, K., et al. (2019). Two distinct interstitial macrophage populations coexist across tissues in specific subtissular niches. Science (80-.). 363, eaau0964. 10.1126/science.aau0964.

Chu, S.G., Villalba, J.A., Liang, X., Xiong, K., Tsoyi, K., Ith, B., Ayaub, E.A., Tatituri, R. V, Byers, D.E., Hsu, F.-F., et al. (2019). Palmitic Acid–Rich High-Fat Diet Exacerbates Experimental Pulmonary Fibrosis by Modulating Endoplasmic Reticulum Stress. Am. J. Respir. Cell Mol. Biol. 61, 737–746. 10.1165/rcmb.2018-0324OC.

Esparza-Baquer, A., Labiano, I., Sharif, O., Agirre-Lizaso, A., Oakley, F., Rodrigues, P.M., Zhuravleva, E., O’Rourke, C.J., Hijona, E., Jimenez-Agüero, R., et al. (2021). TREM-2 defends the liver against hepatocellular carcinoma through multifactorial protective mechanisms. Gut 70, 1345. 10.1136/gutjnl-2019-319227.

Fabre, T., Barron, A.M.S., Christensen, S.M., Asano, S., Bound, K., Lech, M.P., Wadsworth, M.H., Chen, X., Wang, C., Wang, J., et al. (2023). Identification of a broadly fibrogenic macrophage subset induced by type 3 inflammation. Sci. Immunol. 8, eadd8945. 10.1126/sciimmunol.add8945.

Gu, Z., Eils, R., and Schlesner, M. (2016). Complex heatmaps reveal patterns and correlations in multidimensional genomic data. Bioinformatics 32, 2847–2849. 10.1093/bioinformatics/btw313.

Guilliams, M., De Kleer, I., Henri, S., Post, S., Vanhoutte, L., De Prijck, S., Deswarte, K., Malissen, B., Hammad, H., and Lambrecht, B.N. (2013). Alveolar macrophages develop from fetal monocytes that differentiate into long-lived cells in the first week of life via GM-CSF. J. Exp. Med. 210, 1977–1992. 10.1084/jem.20131199.

Habermann, A.C., Gutierrez, A.J., Bui, L.T., Yahn, S.L., Winters, N.I., Calvi, C.L., Peter, L., Chung, M.-I., Taylor, C.J., Jetter, C., et al. (2020). Single-cell RNA sequencing reveals profibrotic roles of distinct epithelial and mesenchymal lineages in pulmonary fibrosis. Sci. Adv. 6, eaba1972. 10.1126/sciadv.aba1972.

Hao, Y., Hao, S., Andersen-Nissen, E., Mauck, W.M., Zheng, S., Butler, A., Lee, M.J., Wilk, A.J., Darby, C., Zager, M., et al. (2021). Integrated analysis of multimodal single-cell data. Cell 184, 3573–3587.e29. 10.1016/J.CELL.2021.04.048.

Henn, D., Chen, K., Fehlmann, T., Trotsyuk, A.A., Sivaraj, D., Maan, Z.N., Bonham, C.A., Barrera, J.A., Mays, C.J., Greco, A.H., et al. (2023). Xenogeneic skin transplantation promotes angiogenesis and tissue regeneration through activated Trem2+ macrophages. Sci. Adv. 7, eabi4528. 10.1126/sciadv.abi4528.

Hoyer, N., Prior, T.S., Bendstrup, E., Wilcke, T., and Shaker, S.B. (2019). Risk factors for diagnostic delay in idiopathic pulmonary fibrosis. Respir. Res. 20, 103. 10.1186/s12931-019-1076-0.

Hughes, T.K., Wadsworth, M.H., Gierahn, T.M., Do, T., Weiss, D., Andrade, P.R., Ma, F., de Andrade Silva, B.J., Shao, S., Tsoi, L.C., et al. (2020). Second-Strand Synthesis-Based Massively Parallel scRNA-Seq Reveals Cellular States and Molecular Features of Human Inflammatory Skin Pathologies. Immunity 53, 878–894.e7. 10.1016/J.IMMUNI.2020.09.015.

Hutchinson, J.P., Fogarty, A.W., McKeever, T.M., and Hubbard, R.B. (2015). In-Hospital Mortality after Surgical Lung Biopsy for Interstitial Lung Disease in the United States. 2000 to 2011. Am. J. Respir. Crit. Care Med. 193, 1161–1167. 10.1164/rccm.201508-1632OC.

Jaeger, B., Schupp, J.C., Plappert, L., Terwolbeck, O., Artysh, N., Kayser, G., Engelhard, P., Adams, T.S., Zweigerdt, R., Kempf, H., et al. (2022). Airway basal cells show a dedifferentiated KRT17highPhenotype and promote fibrosis in idiopathic pulmonary fibrosis. Nat. Commun. 13, 5637. 10.1038/s41467-022-33193-0.

Jung, S.-H., Hwang, B.-H., Shin, S., Park, E.-H., Park, S.-H., Kim, C.W., Kim, E., Choo, E., Choi, I.J., Swirski, F.K., et al. (2022). Spatiotemporal dynamics of macrophage heterogeneity and a potential function of Trem2hi macrophages in infarcted hearts. Nat. Commun. 13, 4580. 10.1038/s41467-022-32284-2.

Katzenelenbogen, Y., Sheban, F., Yalin, A., Yofe, I., Svetlichnyy, D., Jaitin, D.A., Bornstein, C., Moshe, A., Keren-Shaul, H., Cohen, M., et al. (2020). Coupled scRNA-Seq and Intracellular Protein Activity Reveal an Immunosuppressive Role of TREM2 in Cancer. Cell 182, 872–885.e19. 10.1016/j.cell.2020.06.032.

Kaunisto, J., Salomaa, E.-R., Hodgson, U., Kaarteenaho, R., Kankaanranta, H., Koli, K., Vahlberg, T., and Myllärniemi, M. (2019). Demographics and survival of patients with idiopathic pulmonary fibrosis in the FinnishIPF registry. ERJ Open Res. 5, 170–2018. 10.1183/23120541.00170-2018.

Lederer, D.J., and Martinez, F.J. (2018). Idiopathic pulmonary fibrosis. N. Engl. J. Med. 378, 1811–1823. 10.1056/NEJMra1705751.

Liao, M., Liu, Y., Yuan, J., Wen, Y., Xu, G., Zhao, J., Cheng, L., Li, J., Wang, X., Wang, F., et al. (2020). Single-cell landscape of bronchoalveolar immune cells in patients with COVID-19. Nat. Med. 10.1038/s41591-020-0901-9.

Macosko, E.Z., Basu, A., Satija, R., Nemesh, J., Shekhar, K., Goldman, M., Tirosh, I., Bialas, A.R., Kamitaki, N., Martersteck, E.M., et al. (2015). Highly Parallel Genome-wide Expression Profiling of Individual Cells Using Nanoliter Droplets. Cell 161, 1202–1214. 10.1016/j.cell.2015.05.002.

Martinez, F.J., Chisholm, A., Collard, H.R., Flaherty, K.R., Myers, J., Raghu, G., Walsh, S.L.F., White, E.S., and Richeldi, L. (2017). The diagnosis of idiopathic pulmonary fibrosis: current and future approaches. Lancet Respir. Med. 5, 61–71. 10.1016/S2213-2600(16)30325-3.

Mass, E., Ballesteros, I., Farlik, M., Halbritter, F., Günther, P., Crozet, L., Jacome-Galarza, C.E., Händler, K., Klughammer, J., Kobayashi, Y., et al. (2016). Specification of tissue-resident macrophages during organogenesis. Science (80-.). 353, aaf4238. 10.1126/science.aaf4238.

Mayr, C.H., Simon, L.M., Leuschner, G., Ansari, M., Geyer, P.E., Angelidis, I., Strunz, M., Singh, P., Kneidinger, N., Reichenberger, F., et al. (2020). Integrated Single Cell Analysis of Human Lung Fibrosis Resolves Cellular Origins of Predictive Protein Signatures in Body Fluids. SSRN Electron. J. 10.2139/ssrn.3538700.

Mittal, M., Tiruppathi, C., Nepal, S., Zhao, Y.-Y., Grzych, D., Soni, D., Prockop, D.J., and Malik, A.B. (2016). TNFα-stimulated gene-6 (TSG6) activates macrophage phenotype transition to prevent inflammatory lung injury. Proc. Natl. Acad. Sci. 113, E8151–E8158. 10.1073/pnas.1614935113.

Molgora, M., Esaulova, E., Vermi, W., Hou, J., Chen, Y., Luo, J., Brioschi, S., Bugatti, M., Omodei, A.S., Ricci, B., et al. (2020). TREM2 Modulation Remodels the Tumor Myeloid Landscape Enhancing Anti-PD-1 Immunotherapy. Cell 182, 886–900.e17. 10.1016/j.cell.2020.07.013.

Morse, C., Tabib, T., Sembrat, J., Buschur, K.L., Bittar, H.T., Valenzi, E., Jiang, Y., Kass, D.J., Gibson, K., Chen, W., et al. (2019). Proliferating SPP1/MERTK-expressing macrophages in idiopathic pulmonary fibrosis. Eur. Respir. J. 54, 1802441. 10.1183/13993003.02441-2018.

Mulder, K., Patel, A.A., Kong, W.T., Piot, C., Halitzki, E., Dunsmore, G., Khalilnezhad, S., Irac, S.E., Dubuisson, A., Chevrier, M., et al. (2021). Cross-tissue single-cell landscape of human monocytes and macrophages in health and disease. Immunity 10.1016/j.immuni.2021.07.007.

Park, M.D., Reyes-Torres, I., LeBerichel, J., Hamon, P., LaMarche, N.M., Hegde, S., Belabed, M., Troncoso, L., Grout, J.A., Magen, A., et al. (2023). TREM2 macrophages drive NK cell paucity and dysfunction in lung cancer. Nat. Immunol. 24, 792–801. 10.1038/s41590-023-01475-4.

Pesci, A., Ricchiuti, E., Ruggiero, R., and De Micheli, A. (2010). Bronchoalveolar lavage in idiopathic pulmonary fibrosis: What does it tell us? Respir. Med. 104, S70–S73. 10.1016/j.rmed.2010.03.019.

Raghu, G., Remy-Jardin, M., Myers, J.L., Richeldi, L., Ryerson, C.J., Lederer, D.J., Behr, J., Cottin, V., Danoff, S.K., Morell, F., et al. (2018). Diagnosis of idiopathic pulmonary fibrosis. An official ATS/ERS/JRS/ALAT clinical practice guideline. Am. J. Respir. Crit. Care Med. 198, e44–e68. 10.1164/rccm.201807-1255ST.

Ramachandran, P., Dobie, R., Wilson-Kanamori, J.R., Dora, E.F., Henderson, B.E.P., Luu, N.T., Portman, J.R., Matchett, K.P., Brice, M., Marwick, J.A., et al. (2019). Resolving the fibrotic niche of human liver cirrhosis at single-cell level. Nature 575, 512–518. 10.1038/s41586-019-1631-3.

Reyfman, P.A., Walter, J.M., Joshi, N., Anekalla, K.R., McQuattie-Pimentel, A.C., Chiu, S., Fernandez, R., Akbarpour, M., Chen, C.-I., Ren, Z., et al. (2019). Single-cell transcriptomic analysis of human lung provides insights into the pathobiology of pulmonary fibrosis. Am. J. Respir. Crit. Care Med. 199, 1517–1536. 10.1164/rccm.201712-2410OC.

Schyns, J., Bai, Q., Ruscitti, C., Radermecker, C., De Schepper, S., Chakarov, S., Farnir, F., Pirottin, D., Ginhoux, F., Boeckxstaens, G., et al. (2019). Non-classical tissue monocytes and two functionally distinct populations of interstitial macrophages populate the mouse lung. Nat. Commun. 10, 3964. 10.1038/s41467-019-11843-0.

Sivakumar, P., Thompson, J.R., Ammar, R., Porteous, M., McCoubrey, C., I, E.C.I.I., Ravi, K., Zhang, Y., Luo, Y., Streltsov, D., et al. (2019). RNA sequencing of transplant-stage idiopathic pulmonary fibrosis lung reveals unique pathway regulation. ERJ Open Res. 5, 00117–02019. 10.1183/23120541.00117-2019.

Travaglini, K.J., Nabhan, A.N., Penland, L., Sinha, R., Gillich, A., Sit, R. V., Chang, S., Conley, S.D., Mori, Y., Seita, J., et al. (2020). A molecular cell atlas of the human lung from single-cell RNA sequencing. Nature 10.1038/s41586-020-2922-4.

Ucero, A.C., Bakiri, L., Roediger, B., Suzuki, M., Jimenez, M., Mandal, P., Braghetta, P., Bonaldo, P., Paz-Ares, L., Fustero-Torre, C., et al. (2019). Fra-2–expressing macrophages promote lung fibrosis. J. Clin. Invest. 129, 3293–3309. 10.1172/JCI125366.

Xu, Y., Mizuno, T., Sridharan, A., Du, Y., Guo, M., Tang, J., Wikenheiser-Brokamp, K.A., Perl, A.-K.T., Funari, V.A., Gokey, J.J., et al. (2017). Single-cell RNA sequencing identifies diverse roles of epithelial cells in idiopathic pulmonary fibrosis. JCI Insight 1. 10.1172/jci.insight.90558.

Xue, J., Schmidt, S. V., Sander, J., Draffehn, A., Krebs, W., Quester, I., DeNardo, D., Gohel, T.D., Emde, M., Schmidleithner, L., et al. (2014). Transcriptome-Based Network Analysis Reveals a Spectrum Model of Human Macrophage Activation. Immunity 40, 274–288. 10.1016/J.IMMUNI.2014.01.006.

Yu, Y.-R.A., Hotten, D.F., Malakhau, Y., Volker, E., Ghio, A.J., Noble, P.W., Kraft, M., Hollingsworth, J.W., Gunn, M.D., and Tighe, R.M. (2016). Flow Cytometric Analysis of Myeloid Cells in Human Blood, Bronchoalveolar Lavage, and Lung Tissues. Am. J. Respir. Cell Mol. Biol. 54, 13–24. 10.1165/rcmb.2015-0146OC.

Zhang, F., Ayaub, E.A., Wang, B., Puchulu-Campanella, E., Li, Y.-H., Hettiarachchi, S.U., Lindeman, S.D., Luo, Q., Rout, S., Srinivasarao, M., et al. (2020). Reprogramming of profibrotic macrophages for treatment of bleomycin-induced pulmonary fibrosis. EMBO Mol. Med. 12, e12034. 10.15252/emmm.202012034.

Zhang, P., Cao, L., Zhou, R., Yang, X., and Wu, M. (2019). The lncRNA Neat1 promotes activation of inflammasomes in macrophages. Nat. Commun. 10, 1495. 10.1038/s41467-019-09482-6.

